# Diverse laboratory colonies of *Aedes aegypti* harbor the same adult midgut bacterial microbiome

**DOI:** 10.1101/200659

**Authors:** Laura B. Dickson, Amine Ghozlane, Stevenn Volant, Christiane Bouchier, Laurence Ma, Anubis Vega-Rúa, Isabelle Dusfour, Davy Jiolle, Christophe Paupy, Martin N. Mayanja, Alain Kohl, Julius J. Lutwama, Veasna Duong, Louis Lambrechts

**Affiliations:** Insect-Virus Interactions Group, Department of Genomes and Genetics, Institute Pasteur, CNRS URA 3012, Paris, France; Bioinformatics and Biostatistics Hub, C3BI, USR 3756 CNRS, Institut Pasteur, 12 Paris, CNRS URA 3012, Paris, France; Genomics Facility – Biomics Pole, CITECH, Institut Pasteur, Paris, France; Laboratory of Medical Entomology, Environment and Health Unit, Institute Pasteur de la Guadeloupe, Guadeloupe, France; Vector Control and Adaptation, Institut Pasteur de la Guyane, Vectopole Amazonien Emile Abonnenc, Cayenne, French Guiana; MIVEGEC, IRD, CNRS, Univ. Montpellier, Montpellier, France; Centre International de Recherches Médicales de Franceville, Franceville, Gabon; Department of Arbovirology, Uganda Virus Research Institute, Entebbe, Uganda; MRC-University of Glasgow Centre for Virus Research, Glasgow, United Kingdom; Virology Unit, Institut Pasteur in Cambodia, Phnom Penh, Cambodia

**Keywords:** Mosquito, Microbiota, Vectorial capacity, Metagenomics

## Abstract

**Background:** Host-associated microbes, collectively known as the microbiota, play an important role in the biology of multicellular organisms. In mosquito vectors of human pathogens, the gut bacterial microbiota influences vectorial capacity and has become the subject of intense study. In laboratory studies of vector biology, genetic effects are often inferred from differences between geographically and genetically diverse colonies of mosquitoes that are reared in the same insectary. It is unclear, however, to what extent genetic effects can be confounded by uncontrolled differences in the microbiota composition among mosquito colonies. To address this question, we used 16S metagenomics to compare the midgut bacterial microbiome of six recent laboratory colonies of *Aedes aegypti* representing the geographical range and genetic diversity of the species.

**Results:** We found that the diversity, abundance, and community structure of the midgut bacterial microbiome was remarkably similar among the six different colonies of *Ae. aegypti*, regardless of their geographic origin. We also confirmed the relatively low complexity of bacterial communities inhabiting the mosquito midgut.

**Conclusions:** Our finding that geographically diverse colonies of *Ae. aegypti* reared in the same insectary harbor a similar gut bacterial microbiome supports the conclusion that the gut microbiota of adult mosquitoes is environmentally determined regardless of the host genotype. Thus, uncontrolled differences in microbiota composition are unlikely to represent a significant confounding factor in genetic studies of vector biology.

## Background

The mosquito, *Aedes aegypti*, is the main vector of several medically important arboviruses such as Zika, dengue, chikungunya, and yellow fever viruses worldwide. Dengue viruses alone are responsible for 390 million human infections each year [1]. In the absence of vaccines or specific therapeutics for most arboviruses, controlling mosquito vector populations is the primary disease prevention strategy [2]. With the rise of insecticide resistance, the development of novel entomological interventions is underway [3,4]. Critical to the development of these new vector control methods is an improved understanding of the biology of mosquito vectors such as *Ae. aegypti* [5].

Over the last several decades, research efforts have focused on trying to elucidate the genetic [6–8] and environmental [9–12] factors that contribute to variation in the ability of *Ae. aegypti* to transmit human pathogens. Only in recent years, however, has the importance of the microbiota (i.e., host-associated microbes) emerged in vector biology. The gut bacterial microbiota, in particular, influences multiple aspects of the mosquito’s biology including vector competence [13, 14] and has become a topic of extensive research. Manipulation of the bacterial species present in the mosquito midgut has been shown to both increase or decrease the amounts of dengue virus, chikungunya virus, or *Plasmodium falciparum* [15–19]. The composition of the microbiome (i.e., the collective genomes of the microbiota) found in the midgut of mosquitoes is highly variable and dependent on the environment [20–23] and life stage [24–26].

To identify mosquito genetic components of vectorial capacity, researchers often use genetically diverse colonies of mosquitoes reared in the same environment. Observed differences in the vectorial capacity of genetically diverse laboratory colonies is generally attributed to host genetics, and not to potential differences in the gut microbiota, but it remains poorly understood what role mosquito genetics plays in shaping the gut microbiome and whether subtle differences in the microbiome could confound genetic studies. It was recently shown that the gut microbiota can be disrupted by genetic modification of mosquitoes [27]. In more natural insect systems such as the relationship between aphids, intracellular bacteria, and parasitic wasps, bacterial symbionts and not the aphid genotype drive the specificity of the interactions between the aphid and the parasitic wasp [28–30]. In other insects, it has been demonstrated that the gut microbiota contributes to host genotype by parasite genotype interactions [31], suggesting that differences in the microbiota should be considered as an additional factor when elucidating the host genetic contribution to a specific trait.

In *Ae. aegypti*, previous observations of bacterial taxa specific to certain mosquito lines reared in the same insectary [32, 33] raise the question whether differences in gut microbiota could confound interpretation of phenotypic differences among mosquito colonies. To address this question, we used a targeted metagenomics approach to compare the gut microbiome between six recent colonies of *Ae. aegypti* representing the geographical range and genetic diversity of the species. We performed a comprehensive metagenomics analysis including comparison of bacterial diversity within and between samples as well as identifying bacterial genera that are differentially abundant between colonies. Our results provide empirical evidence that adult *Ae. aegypti* mosquitoes reared in the same insectary harbor a similar gut bacterial microbiome, regardless of their geographicorigin.

## Results

To test if laboratory colonies of natural populations of *Ae. aegypti* differ in the diversity and composition of their gut microbiome, the V5-V6 variable region of the 16S ribosomal RNA gene was sequenced in 16-18 individual adult female midguts from each of six recent colonies of *Ae. aegypti*. The six colonies chosen represent the geographical range and genetic diversity of the species (Figure 1) and have spent from three to ten generations in the laboratory (Table 1). The experimental design included three replicate adult cages per colony and the individual libraries were randomized across two separate sequencing runs. Individual midguts were aseptically dissected from nulliparous, 4- to 6-day-old females that had been allowed to mate and feed on sugar following emergence. Out of the 96 individual gut microbiomes sequenced, 2,679 operational taxonomic units (OTUs) representing 587 different bacterial genera were identified. Rarefaction curves showed that a sufficient number of sequencing reads was achieved to comprehensively characterize the bacterial communities in the midgut (Supplemental Figure 1).

**Figure 1:**
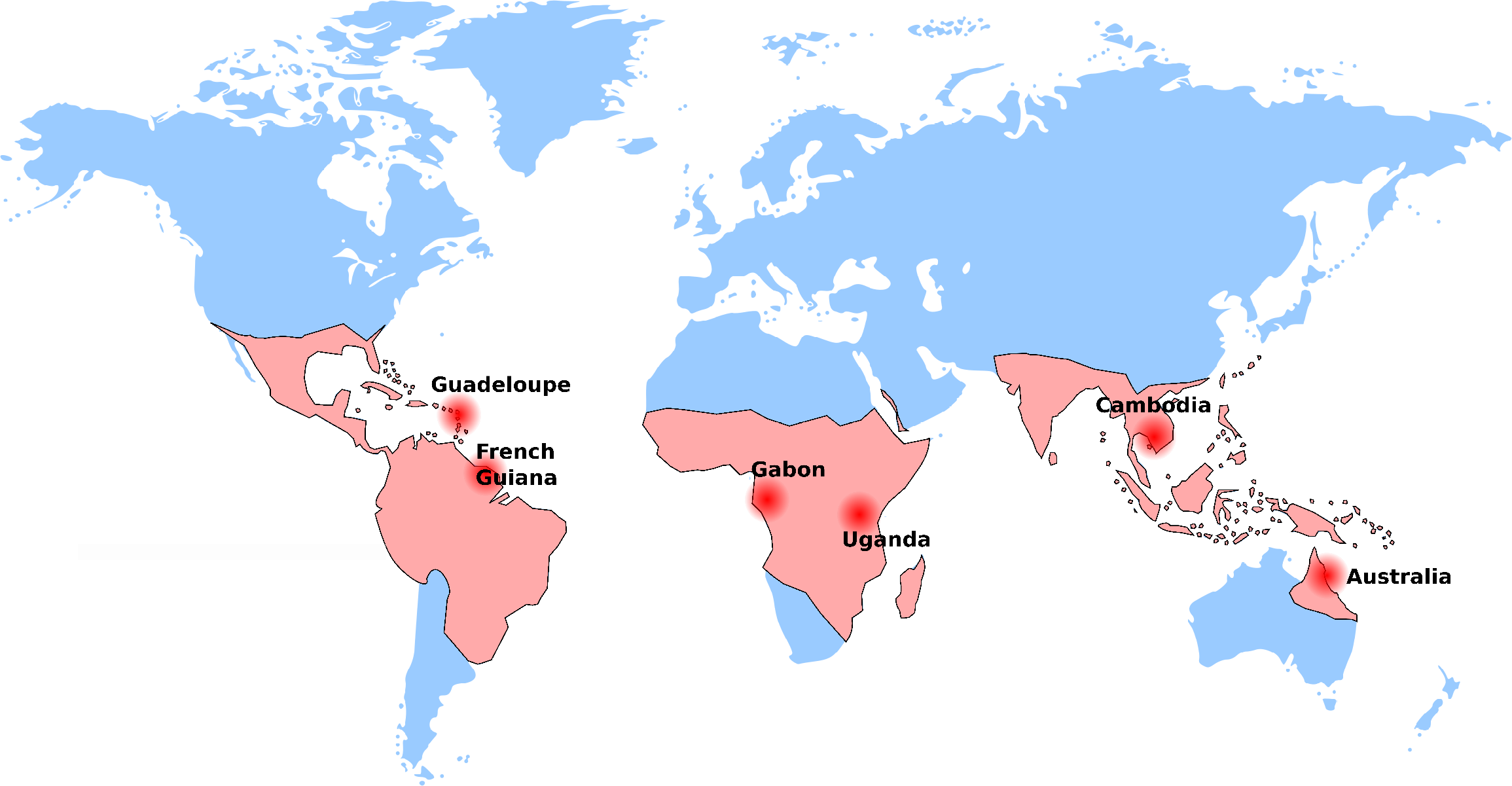
World map showing the origin of the *Ae. aegypti* colonies used in the study overlaid with the approximate global distribution of *Ae. aegypti* adapted from Kraemer et al. [47, 48]. The colonies were initiated on different years and represent different generation times in the laboratory (Table 1).

**Table 1.**
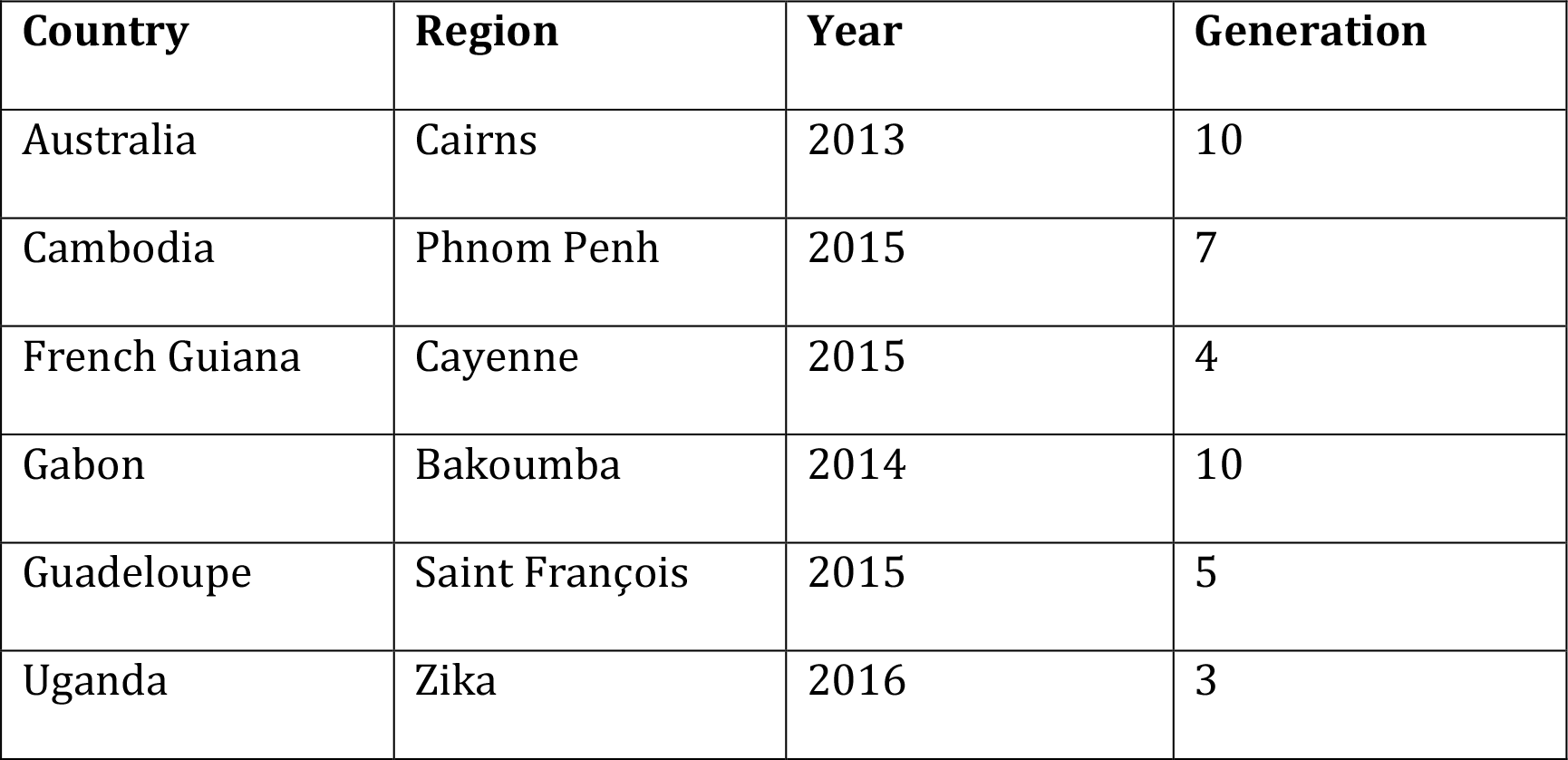
*Aedes aegypti* colonies included in this study. The country and region of origin, year of collection, and number of generations spent in the laboratory prior to the study are shown.

| Country | Region | Year | Generation |
| --- | --- | --- | --- |
| Australia | Cairns | 2013 | 10 |
| Cambodia | Phnom Penh | 2015 | 7 |
| French Guiana | Cayenne | 2015 | 4 |
| Gabon | Bakoumba | 2014 | 10 |
| Guadeloupe | Saint François | 2015 | 5 |
| Uganda | Zika | 2016 | 3 |

To determine if the gut microbiome of each *Ae. aegypti* colony varies in the diversity of bacterial species present, the within-colony diversity was evaluated by determining the genus richness and the Shannon diversity index. No differences in the levels of richness (Figure 2A) or in Shannon diversity index (Figure 2B) were observed between the colonies (ANOVA: F = 1.125, *p* value = 0.353 and F = 0.522, *p* value = 0.759, respectively). In addition, the taxonomical abundance of bacteria was highly similar between the colonies, indicating the dominant bacterial genera in the midgut are not dependent on the colony (Figure 3).

**Figure 2:**
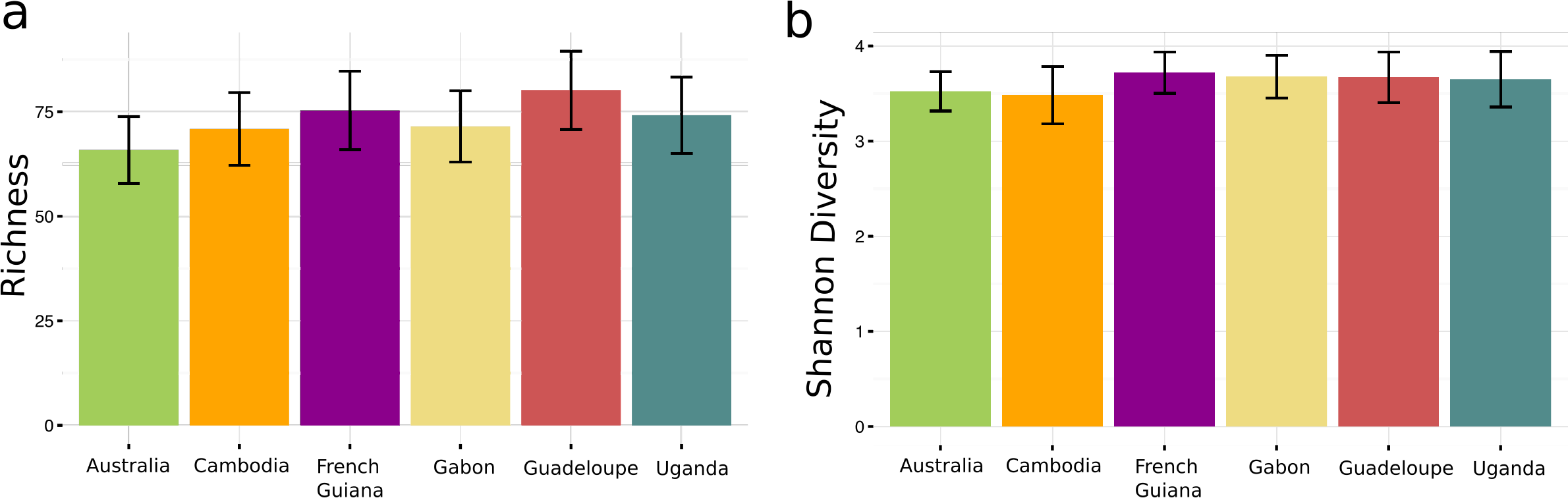
Genetic diversity of the gut bacterial communities is similar between diverse colonies of *Ae. aegypti*. The genus richness (a) and Shannon diversity index (b) were calculated for each colony representing 16-18 individual midguts from 3 replicate cages dissected 4-6 days after adult emergence. Genus richness is the number of bacterial genera identified in each colony. The Shannon diversity index accounts for the relative abundance of each bacterial genus. Error bars represent 95% confidence intervals. No difference in richness (ANOVA: F = 1.125, *p* value = 0.353) or in Shannon index (ANOVA: F = 0.522, *p* value = 0.759) was detected between colonies.

**Figure 3:**
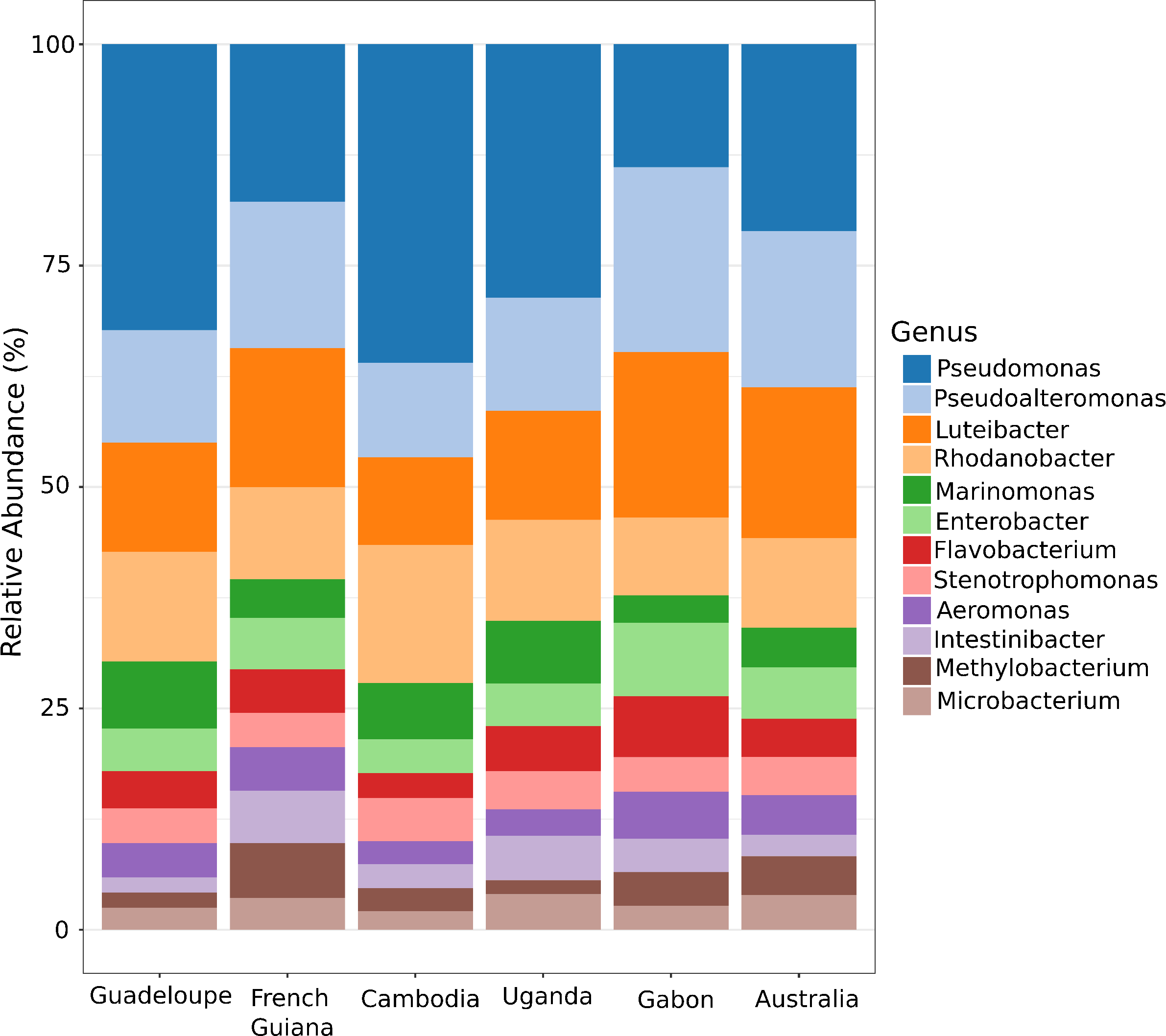
The dominant bacterial genera found in the midgut are similar among diverse colonies of *Ae. aegypti*. The abundance of the 12 most abundant genera is shown for each colony representing 16-18 individual midguts from 3 replicate cages dissected 4-6 days after adult emergence. Bacterial genera were assigned to OTUs clustered with a 97% cutoff using the SILVA database (https://www.arb-silva.de).

To identify dissimilarities in the bacterial community structure between the gut microbiome of laboratory colonies of *Ae. aegypti*, principal coordinates analysis (PCoA) was performed based on a Bray-Curtis dissimilarity matrix. The PCoA showed that the bacterial community structures of all six colonies were highly similar to each other (*p* value = 0.752) (Figure 4A). In addition, no differences in the bacterial community structure were observed between the replicate cages of each colony, however the bacterial community structure differed between sequencing runs (Supplemental Figure 2). The reason for the run effect is unclear but it could reflect preferential clustering of specific sequences on the flow cell. Although the community structure of gut microbiome of the colonies was similar overall, we tested whether some specific bacterial taxa were differentially abundant. Out of the 587 bacterial genera identified, only zero to six genera were differentially abundant in pairwise comparisons of the six colonies (Supplemental Table 1; Figure 4B) resulting in 98-100% similarity in the abundance of genera present between colonies.

**Figure 4:**
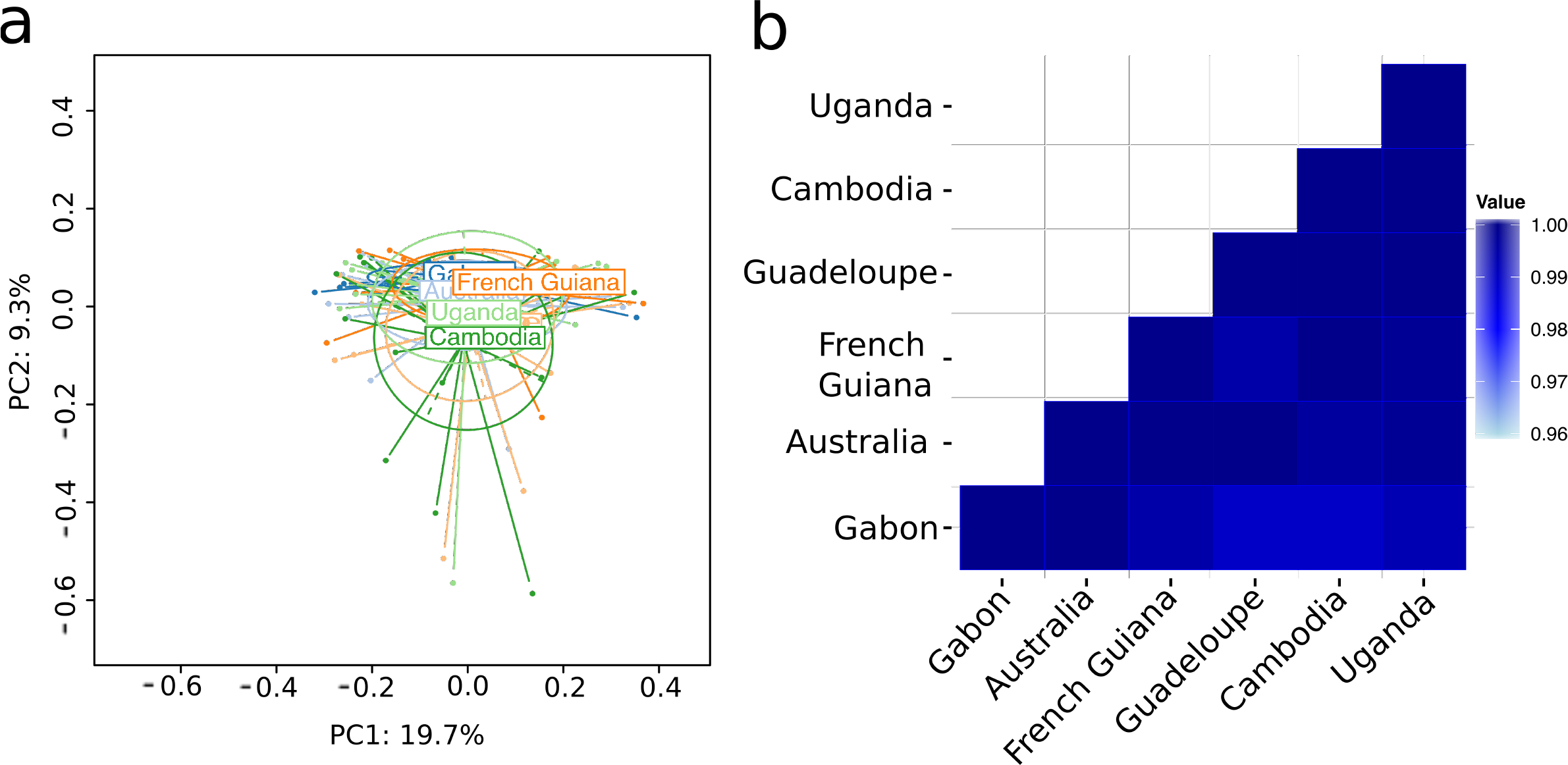
The midgut bacterial community structure is similar between diverse colonies of *Ae. aegypti*. Bacterial community structures between colonies are compared by (a) principal coordinates analysis (PCoA) and (b) pairwise differential abundance analysis. PCoA is based on a Bray-Curtis dissimilarity matrix and indicates a lack of overall differences (PERMANOVA: *p* value = 0.752). Results of differential abundance analysis are shown for each pair of colonies as the proportion of all bacterial genera identified (n=587) that were non-significantly differentially abundant after correction for multiple testing.

## Discussion

We performed a 16S metagenomics analysis to compare the midgut microbiome of six recent colonies of *Ae. aegypti* reared in the same insectary environment. The six colonies were chosen to represent the natural global distribution ofthe species. Although these colonies represent different genetic backgrounds and different generation times in the laboratory, the gut microbiome was highly similar among all six colonies. We did not observe any differences in the diversity of the bacterial communities or in the bacterial community structure within the gut. The taxonomical abundance was also similar between the colonies with 98-100% identity in the abundance of bacterial genera present between colonies. The data also confirmed the relatively low complexity of bacterial communities typically found in the gut of insects [34–35].

Other studies that have compared the midgut microbiome of various laboratory colonies of *Ae. aegypti* observed differences in the taxonomical identification of specific bacterial species [32, 33]. Although these studies reported differences in the abundance of specific taxa between colonies of *Ae. aegypti*, no difference in the bacterial community structure was in fact observed. Furthermore, the colonies tested in previous studies have been maintained in the laboratory for five to 80 years before their microbiome was examined. It is possible that large differences in the number of generations spent in the laboratory between these studies and ours, resulted in our different observations. Possibly, preferential associations between mosquito genotypes and specific laboratory bacteria may evolve over a long colonization history. This hypothesis remains to be tested.

Researchers often use genetically diverse colonies of mosquitoes reared in the same environment to identify mosquito genetic components of vectorial capacity. In such studies, differences in microbiota could confound interpretation of phenotypic differences among mosquito colonies. The present study does not support this hypothesis in the case of *Ae. aegypti*. While this may be the case in some insect systems [28–31], our study provides evidence that the midgut microbiome of colonized *Ae. aegypti* is highly similar and most likely will not confound genetic studies of vector biology.

Although we did not genotype the *Ae. aegypti* colonies used in this study, it is well accepted that *Ae. aegypti* from sub-Saharan Africa belong to a different phylogenetic cluster than pan-tropical *Ae. aegypti* from elsewhere in the world [36–40]. At a more local scale, populations of *Ae. aegypti* sampled from distinct locations are usually genetically distinct [41–43]. Accordingly, we assume that the colonies that we tested in fact represent various genotypes of *Ae. aegypti*. We conclude that mosquito genotype does not influence the microbiome of laboratory-bred *Ae. aegypti*, further demonstrating the importance that the environment plays in shaping the gut microbiome of *Ae. aegypti*. However, this may not be the case in a more natural system. One can imagine that withina given environment, the mosquito genotype may influence the composition of the midgut microbiome and this should be explored further.

Alimitation of our study was that we only dissected midguts at one time point. Recent results from Short et al. [33] suggest that differences between colonies may exist at different times following adult emergence. It is possible that differences in the gut microbiome between our colonies would have been observed if we had sampled the midguts sooner or later after adult emergence. Since our primary goal was to determine how the gut microbiome of our colonies impacted studies of vector competence, we chose a time point after adult emergence that related to the time when an infectious blood meal is usually offered in vector competence assays.

One potentially important implication of our results is that the same mosquito strain reared in different laboratories might display different phenotypes due to a different gut microbiome. We found that the gut microbiome of mosquito colonies was entirely determined by the insectary environment regardless of the mosquito genotype. It follows that the same mosquito strain exposed to a different environment could host a different gut microbiota. This could undermine the relevance of reference strains that are shared by different laboratories. It will be interesting in future studies to compare the gut bacterial microbiome of the same mosquito strain reared in different insectaries.

## Conclusions

Our finding that geographically diverse colonies of *Ae. aegypti* reared in the same insectary harbor a similar gut bacterial microbiome supports the conclusion that the gut microbiota of adult mosquitoes is environmentally determined, regardless of the host genotype. Thus, uncontrolled differences in microbiota composition are unlikely to represent a significant confounding factor in genetic studies of vector biology.

## Methods

### Mosquito colonies and sample preparation

Six *Ae. aegypti* colonies were chosen to represent the worldwide distribution of the species (Figure 1;Table 1). Eggs from each of these colonies were simultaneously hatched in dechlorinated tap water under reduced air pressure for one hour and 200 first–instar larvae from each colony were sorted into 24 x 34 x 9 cm plastic trays. The larvae were fed on a standard diet of Tetramin fish food (Tetra) every other day until pupation. Immediately following emergence, adults (males and females) were randomly separated into three replicate cages per mosquito colony. They were maintained under standard insectary conditions(28°C, 70% relative humidity and 12h light:12h dark cycle) for 4-6 days and allowed to mate and feed on sugar.

Midguts were dissected from adult females under sterile conditions in a biosafety cabinet. Each mosquito was surface sterilized in 70% ethanol for 3-5 minutes and washed three times in sterile 1x phosphate-buffered saline (PBS). Midguts were dissected in a drop of sterile 1x PBS and DNA from individual midguts was extracted as previously reported [9]. Briefly, individual midguts were ground in 300 μl of 20 mg/ml lysozyme dissolved in Qiagen ATL buffer in a sterile tube containing grinding beads. The samples were homogenized for two rounds of 30 seconds at 6,700 RPM (Precellys 24, Bertin Technologies) and DNA was extracted following the Qiagen DNeasy recommended pre-treatment protocol for Gram-positive bacterial samples. To control for contamination of bacteria introduced during the midgut dissections, DNA extractions, and PCR steps, negative controls were made by extracting DNA from blank 1x PBS that was used during the washing steps and by performing negative PCR reactions.

### 16S sequencing

Custom-made PCR primers were designed to amplify the hypervariable V5-V6 region of the bacterial 16S ribosomal RNA gene from midguts as previously described [9]. Purified DNA from each midgut sample was amplified in triplicate by 40 cycles of PCR using Expand High-Fidelity polymerase (Sigma-Aldrich) following manufacturer instructions. To improve PCR sensitivity, 0.15 μl T4gene32 and 0.5 μl 20mg/ml bovine serum albumin (BSA) were added per reaction with6 ml of template DNA. The three PCR reactions were pooled and the PCR products purified using Agencourt AMPure XP magnetic beads (Beckman Coulter). The purified PCR products were quantified by Quant-iT PicoGreen dsDNA fluorometric quantification (ThermoFisher Scientific) and pooled for sequencing in paired-end on the Illumina MiSeq platform using the 500-cycle v2 chemistry (Illumina). On average, 16-18 individual midguts (6 individuals per adult replicate cage) were sequenced per *Ae. aegypti* colony. In order to achieve enough reads per sample, the sequencing was done in two separate runs. Libraries from each colony and each replicate were dispersed evenly between the two sequencing runs. Five libraries were removed from further analysis due to a low number of reads. Raw sequences were deposited to the European Nucleotide Archive under accession number PRJEB22905.

## Data analysis

To account for possible contamination at various steps in the sample-processing pipeline, the sequencing reads were corrected with the reads from the negative controls. The sequencing reads from each sample were mapped to the reads found in the negative controls using Bowtie2 [44]. Reads that mapped to reads in the negative controls were removed from the analysis. Read filtering, OTU clustering and annotation were performed with the MASQUE pipeline. (https://github.com/aghozlane/masque) as previously described [45]. A total of 2,679 OTUs were obtained at 97% sequence identity threshold. Genus richness and Shannon diversity index were compared by analysis of variance (ANOVA). All other statistical analyses were performed with SHAMAN (shaman.c3bi.pasteur.fr) as previously described [9]. Briefly, the normalization of OTU counts was performed at the OTU level using the DESeq2 normalization method. After normalization, an additional six individuals were removed due to low size factors. In SHAMAN, a generalized linear model (GLM) was fitted and vectors of contrasts were defined to determine the significance in abundance variation between sample types. The GLM included the main effect of the *Ae. aegypti* colony, the main effect of replicate cage, the main effect of sequencing run and the interaction between colony and replicate. The resulting *p* values were adjusted for multiple testing according to the Benjamini and Hochberg procedure [46]. Principal coordinates analysis (PCoA) was performed with the ade4 R package (v1.7.6) using a Bray-Curtis dissimilarity matrix. Permutational multivariate analysis of variance (PERMANOVA)was performed in the vegan R package (v2.4.3) as a distance-based method to test the statistical significance of the association between bacterial community structureand mosquito colony.

## Authors’ contributions

LBD and LL designed the study. AVR, ID, DJ, CP, MNM, AK, JJL, and VD provided mosquito specimens to initiate colonies. LBD carried out the experiments. LBD, CB, and LM performed the sequencing. LBD, AG, SV, and LL analyzed the data. LBD and LL wrote the manuscript. All authors read and approved the final manuscript.

## Acknowledgements

We are grateful to Gordana Rašić and Ary Hoffmann for providing the mosquito colony from Cairns and to Borin Peng for the mosquito collection in Phnom Penh. We thank Claire Valiente Moro,Guillaume Minard and the Lambrechts lab members for their insights.

---

**Supplemental Table 1:** Identification of bacterial genera that are differentially abundant in pairwise comparisons of colonies. The lack of a comparison between two colonies indicates that no bacterial genera were significantly different between them.

**Supplemental Figure 1:** Rarefaction curves for the individual samples used in the analysis at the genera level. The curves show the number of detected bacterial genera as a function of the number of reads analyzed per sequencing library. Each curve represents a single midgut sample.

**Supplemental Figure 2:** The midgut bacterial communities are highly structured by sequencing run. The cluster dendrogram of individual midgut samples based on a Bray-Curtis dissimilarity matrix shows that sequencing run, and not the identity of the mosquito colony, determines bacterial community relatedness. Midgut samples are represented by numbers color coded by sequencing run. Dark blue samples were sequenced in the first run, whereas light blue samples were sequenced in the second run.

